# Targeted NAD^+^ Delivery for Intimal Hyperplasia and Re-endothelialization: A Novel Anti-restenotic Therapy Approach

**DOI:** 10.1101/2024.02.20.581249

**Authors:** Li Yin, Yao Tong, Zain Husain Islam, Kaijie Zhang, Ruosen Xie, Jacobus Burger, Nicholas Hoyt, Eric William Kent, William Aaron Marcum, Campbell Johnston, Rohan Kanchetty, Zoe Tetz, Sophia Stanisic, Yitao Huang, Lian-Wang Guo, Shaoqin Gong, Bowen Wang

## Abstract

Endovascular interventions often fail due to restenosis, primarily caused by smooth muscle cell (SMC) proliferation, leading to intimal hyperplasia (IH). Current strategies to prevent restenosis are far from perfect and impose significant collateral damage on the fragile endothelial cell (EC), causing profound thrombotic risks. Nicotinamide adenine dinucleotide (NAD^+^) is a co-enzyme and signaling substrate implicated in redox and metabolic homeostasis, with a pleiotropic role in protecting against cardiovascular diseases. However, a functional link between NAD^+^ repletion and the delicate duo of IH and EC regeneration has yet to be established. NAD^+^ repletion has been historically challenging due to its poor cellular uptake and low bioavailability. We have recently invented the first nanocarrier that enables direct intracellular delivery of NAD^+^ *in vivo*. Combining the merits of this prototypic NAD^+^-loaded calcium phosphate (CaP) nanoparticle (NP) and biomimetic surface functionalization, we created a biomimetic P-NAD^+^-NP with platelet membrane coating, which enabled an injectable modality that targets IH with excellent biocompatibility. Using human cell primary culture, we demonstrated the benefits of NP-assisted NAD^+^ repletion in selectively inhibiting the excessive proliferation of aortic SMC, while differentially protecting aortic EC from apoptosis. Moreover, in a rat balloon angioplasty model, a single-dose treatment with intravenously injected P-NAD^+^-NP immediately post angioplasty not only mitigated IH, but also accelerated the regeneration of EC (re-endothelialization) *in vivo* in comparison to control groups (i.e., saline, free NAD^+^ solution, empty CaP-NP). Collectively, our current study provides proof-of-concept evidence supporting the role of targeted NAD^+^ repletion nanotherapy in managing restenosis and improving re-endothelialization.

## 1. Introduction

The advent of endovascular interventions such as balloon angioplasty and stenting has ushered in a new era for the combat against cardiovascular and peripheral vascular diseases. However, the long-term patency of the reconstructed vessels has been consistently compromised, primarily caused by restenosis – the re-narrowing or re-occlusion of the target vessels.^1,2^ Over the past two decades, several generations of drug-eluting stents and drug-coated balloons have been introduced to specifically address the persistent recurrence of restenosis^3,4^. Through device-enabled intraluminal delivery of often toxic drugs (e.g., paclitaxel and sirolimus), anti-restenotic effects could be achieved via inhibiting the excessive proliferation of vascular smooth muscle cell (SMC), or intimal hyperplasia (IH).^5,6^ In comparison to bare metal stents and plain balloon angioplasty, these drug-eluting/-coated devices are associated with reduced restenotic rates and hence improved post-intervention outcomes.^7^

Unfortunately, emerging evidence suggests that drug-eluting/-coated devices create excessive damage to the fragile endothelial cell (EC), leading to impaired re-endothelialization and hence increased risk of thrombosis and related adverse events.^8-10^ The use of dual-antiplatelet therapy following endovascular interventions is required to reduce the thrombotic risk, yet its extended use not only increases the risk of bleeding but also incurs significant financial burdens.^11^ Indeed, while the anti-restenotic drugs successfully block SMC proliferation and IH, they simultaneously exacerbate the apoptotic and pro-inflammatory responses of the adjacent ECs.^12-14^ A growing body of literature, including ours, confirms that arterial/aortic ECs are far more sensitive to the cytotoxicity incurred by commonly used anti-restenotic agents than SMCs.^14-17^ Therefore, it remains a paramount challenge to identify novel intervention targets and therapeutic agents that can selectively block the excessive proliferation of SMC and IH, while preserving or even rescuing the integrity of EC following endovascular intervention.

Recently, nicotinamide adenine dinucleotide (NAD^+^) supplementation has been widely proven to be a viable strategy for cardiovascular health.^18^ NAD^+^ is an important endogenous co-enzyme that critically contributes to redox and nutrient metabolic homeostasis. It also serves as signaling substrates for various key pathways with diverse physiological consequences in the vascular system.^19^ Disturbed NAD^+^ homeostasis and decline of intracellular NAD^+^ pool have been widely associated with vascular aging, hypertension, and atherosclerosis.^20-22^ Serendipitously, our very own study as well as others demonstrate a pivotal role of NAD^+^ repletion in protecting EC from sepsis-induced disease.^23^ The role of NAD^+^ in SMC, despite sparsely documented, is suggested to reinstate SMC phenotypic integrity, based on evidence derived from rodent models of aortic aneurysms.^24^ Collectively, NAD^+^ repletion may constitute a viable strategy for EC-protective, yet SMC-inhibitive management of IH, which has not been achieved in current endovascular interventions.

Herein, we aim to determine the pre-clinical outcomes of targeted NAD^+^ repletion therapy in terms of both promoting re-endothelialization and inhibiting IH. This new therapy is enabled by the concerted innovations from two fronts: direct intracellular NAD^+^ repletion, and IH-targeting nanomedicine. On the one hand, our team recently developed the first-of-its-kind calcium phosphate nanoparticle (i.e., NAD^+^-NP) that demonstrated efficient NAD^+^ repletion in murine models of sepsis. Prior attempts in direct NAD^+^ delivery *in vivo* all failed, primarily due to the low bioavailability and lack of direct cellular uptake of NAD^+^. On the other hand, our team and others pioneered the concept of nanoparticle (NP) biomimetic surface functionalization with cell membrane vesicles, such as platelets, which could grant NPs with IH-targeting capacity and biocompatibility.^25^ The combined innovation allowed us, for the first time, to probe targeted NAD^+^ repletion as a new avenue in achieving differential management of the two seemingly opposite objectives: anti-restenosis (inhibiting SMC proliferation) and re-endothelialization (promoting EC regeneration).

## 2. Methods and Materials

### 2.1 Materials

β-nicotinamide adenine dinucleotide (NAD+), calcium chloride (CaCl2), and IGEPAL CO-520 were purchased from Sigma-Aldrich (St. Louis, MO, USA). Disodium hydrogen phosphate (Na2HPO4) was purchased from Dot Scientific Inc., (Burton, MI, USA). Dioleoyhosphatydic acid (DOPA) and L-α-phosphatidylcholine (Soy PC) were obtained from Avanti Polar Lipids (Alabaster, AL, USA). ATTO550-conjugated oligoRNA was bought from Integrated DNA Technologies (Coralville, IA, USA). Cyclohexane and chloroform were purchased from Thermo Fisher Scientific (Fitchburg, WI, USA).

### 2.2 Preparation of biomimetic platelet membrane-coated and NAD^+^-loaded calcium phosphate NP (P-NAD^+^-NP)

NAD^+^-NP was synthesized utilizing a water-in-oil microemulsion method, followed by thin-film hydration, as described in our previous publication^23^. Briefly, two reverse water-in-oil microemulsions, each of 25 ml, were prepared in a cyclohexane/IGEPAL CO-520 mixture with a volume ratio of 71:29. The first emulsion contained a 2.5 M CaCl2 solution (500 μl) and 1 mg of NAD+. The second emulsion comprised a 25 mM Na2HPO4 solution (500 μl, pH 9) with 3 mg of DOPA. The two emulsions were mixed and stirred for 30 minutes. The emulsion was then broken by adding 50 ml of ethanol and then the mixture was centrifuged at 12000 g to remove the oil phase. The pellet was sequentially washed twice with ethanol and once with 70% ethanol. Subsequently, the nanoparticles were redispersed in chloroform containing Soy PC (3 mg) and cholesterol (0.3 mg.) Following the removal of chloroform using a rotary evaporator, the resultant lipid film was rehydrated with 500 μl Tris-HCl buffer (10 mM, pH 7.4) to produce the NAD^+^-NP.

Biomimetic nanoparticles were prepared by coating the platelet membrane on the surface of the NAD^+^-NP via the extrusion method^26,27^. In short, platelet membrane vesicles were mixed with NAD^+^-NP in a 1:1 weight ratio of membrane protein to NAD^+^-NP. This mixture was then sequentially extruded through polycarbonate membranes with pore sizes of 400 nm followed by 200 nm using an Avanti mini extruder. Sterile conditions were maintained throughout the process. The empty biomimetic NP was prepared following the same procedure by extruding empty NP (without encapsulating NAD^+^) with the platelet membrane vesicles. In the biodistribution study, ATTO550-labeled biomimetic nanoparticles were prepared by co-encapsulating 100 μg ATTO550-labeled oligoRNA and 1 mg NAD^+^.

### 2.3 Characterization of biomimetic P-NAD^+^-NP

The hydrodynamic diameter and zeta potential of P-NAD^+^-NP and related nanoparticles were measured by dynamic light scattering (DLS) using a ZetaSizer Nano ZS90 spectrometer (Malvern Instruments, USA). The morphology of these nanoparticles was determined by transmission electron microscopy (TEM, FEI Tecnai G2 F30 TWIN 300 KV, E.A. Fischione Instruments, Inc., USA). The loading efficiency and loading content of NAD^+^ in the nanoparticles were measured using high-performance liquid chromatography (HPLC, Hitachi Elite LaChrom, USA) with absorbance measured at 260 nm^28^. For the stability study, freshly made P-NAD^+^-NP were stored in 10 mM Tris HCl buffer (pH 7.4) at 4°C and their hydrodynamic diameters were measured daily by DLS.

### 2.4 Animal protocol

All animal studies conform to the National Institutes of Health Guide for the Care and Use of Laboratory Animals and The ARRIVE guidelines (Animal Research: Reporting of *In Vivo* Experiments). Animal research protocol was approved by the Animal Care and Use Committee (ACUC) at the University of Virginia.

### 2.5 Balloon angioplasty procedure in rat carotid arteries

Balloon Angioplasty for common carotid arteries was performed in male Sprague-Dawley rats (350g-380g) as described previously^29^. In brief, the rats were anesthetized with isoflurane (5% for induction and 2-3% for maintenance). After preparing the skin, a midline neck incision was made to expose the left common carotid artery. The branches of the artery were looped to avoid reflux of blood. To injure the artery, we inserted a 2-F Fogarty arterial embolectomy catheter (Edward Life Science, Irvine, CA) through an arteriotomy on the external artery. The balloon catheter was inflated at 1.5 atm and then withdrawn until the carotid bifurcation. After three times repeating this action followed by a fourth cycle with concomitant rotation of the catheter during withdrawal, the catheter was removed. The external carotid artery was ligated permanently, and the blood flow was resumed. Throughout the procedure, the rat was kept anesthetized via isoflurane, inhaling at a flow rate of 2-4L/min. Carprofen (5mg/kg) and Bupivacaine were subcutaneously injected.

Right after the procedure, we provided a single-dose regimen via intravenous injection with the following treatments: saline, empty biomimetic CaP-NP (empty NP), free NAD^+^ solution (free NAD^+^), and biomimetic P-NAD^+^-NP – all at a dose equivalent to 10 mg/kg NAD^+^ payload.

The common carotid arteries were collected two weeks after the procedure. During the harvest, the animals were anesthetized with isoflurane and the arteries were collected following perfusion fixation at a physiological pressure of 100mmHg. Animals were euthanized in a chamber gradually filled with CO2.

### 2.6 Evans blue staining for re-endothelialization evaluation

Balloon Angioplasty for common carotid arteries was performed in the same way as described above. Seven days after the procedure, re-endothelialization of the arteries was evaluated using an Evans blue staining assay as previously reported^29^. One mL of 0.5% Evans blue solution (Sigma-Aldrich, St. Louis, MO) was injected through the tail vein. After 30 mins of injection, the rat was anesthetized as described above. The common carotid artery was collected following saline and RNAlater perfusion. Animals were euthanized in a chamber gradually filled with CO2.

### 2.7 Histological and morphometric analysis

Intimal hyperplasia: Paraffin cross sections (5 μm thick) were cut at equally spaced intervals using a microtome (Leica) and then stained (Hematoxylin and Eosin, abbreviated as H&E; and Masson Trichrome, abbreviated as MT) for morphometric analysis of intimal hyperplasia, as described previously. Morphometric parameters were measured on the sections and calculated by using ImageJ software: lumen area, the area inside internal elastic lamina (IEL area), the area inside external elastic lamina (EEL area), intima area (= IEL area – lumen area), and media area (=EEL area – IEL area). Intimal hyperplasia (IH), the ratio of intima area versus media area (I/M) was calculated by (IEL area – lumen area) / (EEL area – IEL area). 3-6 sections from each rat were measured, and the measurement was pooled from all sections to generate a mean value for each animal. The mean values from all the animals in each treatment group were averaged, and then the standard error of the mean (SEM) was calculated.

Re-endothelialization evaluation: The common arteries were longitudinally opened for image acquisition. Re-reendothelialization area was measured and calculated using ImageJ software: the blue staining area (no endothelium coverage) and the no-staining area (with endothelium coverage). Base level endothelium coverage was determined by arteries collected immediately after the procedure and was subtracted from Day 7 values. Re-reendothelialization (%) = (No-staining area – baseline) / (Blue staining area+ No-staining area – baseline).

All the measurements were performed by an independent researcher who was blinded to the experimental treatment.

### 2.8 Cell culture

Primary culture of human aortic SMC (HASMC), human aortic EC (HAEC), and their respective optimal culture media (SmBm2/EBM2 basal medium for experiments, and SmGm2/EGM2 complete medium for expansion) were purchased from Lonza (Walkersville, MD). Primary cells between passages 4-8 were cultured at 37°C with 5% CO2 as previously described.30,31 For subculture, Trypsin-EDTA and Accutase (Thermo Fisher Scientific, Waltham, MA) were used for cell detachment of HASMC and HAEC, respectively, as established in our prior reports.^30,31^

### 2.9 mRNA Extraction and real-time quantitative polymerase chain reaction (qPCR)

mRNA from the primary culture of human aortic SMC and EC, as well as rat carotid artery tissue homogenates, was isolated using TRIzol according to the manufacturer’s manual. Following quantitation using NanoDrop, 1 μg of mRNA was used for the synthesis of first-strand cDNA using the High-Capacity cDNA Reverse Transcription Kit (ThermoFisher Scientific, Waltham, MA). cDNA templates were then amplified in triplicates with SYBR Green PCR Master Mix using the QuantStudio3 real-time qPCR system (Applied Biosystems, Carlsbad, CA). The sequences for each primer set are provided in the Supplementary Document (Table S1).

### 2.10 Statistical Analysis

Statistical analysis was performed using GraphPad Prism 9 and 10. Prior to parametric analysis, the normality and equal variance of data sets were tested. For two-group comparisons, a Student’s *t*-test was performed. For multiple group-wise comparisons, a One-way analysis of variance (ANOVA) was performed followed by appropriate post-hoc analysis.

## 3. Results

### 3.1 Fabrication and Characterization of Biomimetic P-NAD^+^-NP

NAD^+^-NP was prepared by a water-in-oil reverse microemulsion process followed by thin-film hydration, resulting in nanoparticles with an average hydrodynamic diameter of approximately 165 nm (Figure 1A). The morphology of NAD^+^-NP was determined by TEM (Supplementary Figure 1). To fabricate the platelet membrane-coated NAD^+^-NP (P-NAD^+^-NP), an extrusion technique was applied. The optimal weight ratio for platelet membrane/NAD^+^-NP was first determined to be 1:1 (Supplementary Figure 2). Compared to NAD^+^-NP, the average hydrodynamic diameter of the P-NAD^+^-NP increased to about 190 nm, with the increment attributable to the platelet membrane coating (Figure 1A). The morphology of P-NAD^+^-NP was confirmed by TEM, revealing a diameter of about 170 nm (Figure 1B). The successful coating of the platelet membrane was further evidenced by the alteration in zeta potential, shifting from -5 mV to about -30 mV, which is comparable to the surface charge of platelet membrane^32^ (Figure 1C). The loading efficiency of NAD^+^ within the P-NAD^+^-NP was determined to be 55% by HPLC, corresponding to a loading content of 5.9%. Stability assessments demonstrated that P-NAD^+^-NP remained stable for at least one week when stored at 4°C (Figure 1D).

**Figure 1.**
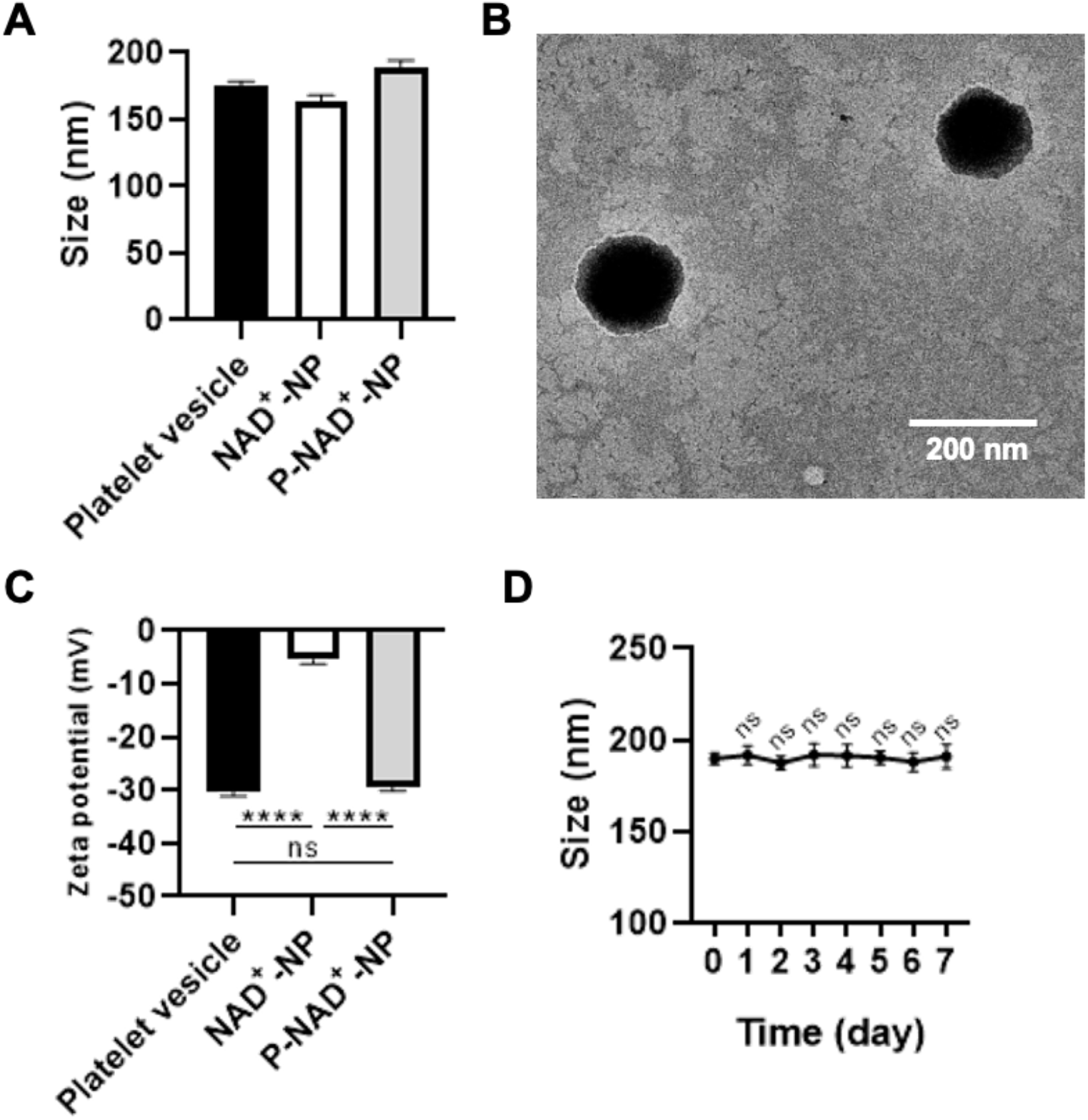
Characterization of biomimetic platelet membrane-coated and NAD^+^-loaded CaP-NP (P-NAD^+^-NP) (A) Hydrodynamic diameters of platelet vesicle, NAD^+^-NP and P-NAD^+^-NP characterized by DLS. (B) Morphology of P-NAD^+^-NP characterized by transmission electron microscopy. (C) Zeta potentials of platelet vesicles, non-biomimetic NAD^+^-NP, and the biomimetic P-NAD^+^-NP measured by dynamic light scattering. (D) Hydrodynamic diameters of P-NAD^+^-NP stored at 4°C for one week by DLS. Presented as Mean±SEM. \*\*\*\**P* < 0.0001. One-way ANOVA with Tukey post-hoc analysis.

**Figure 2.**
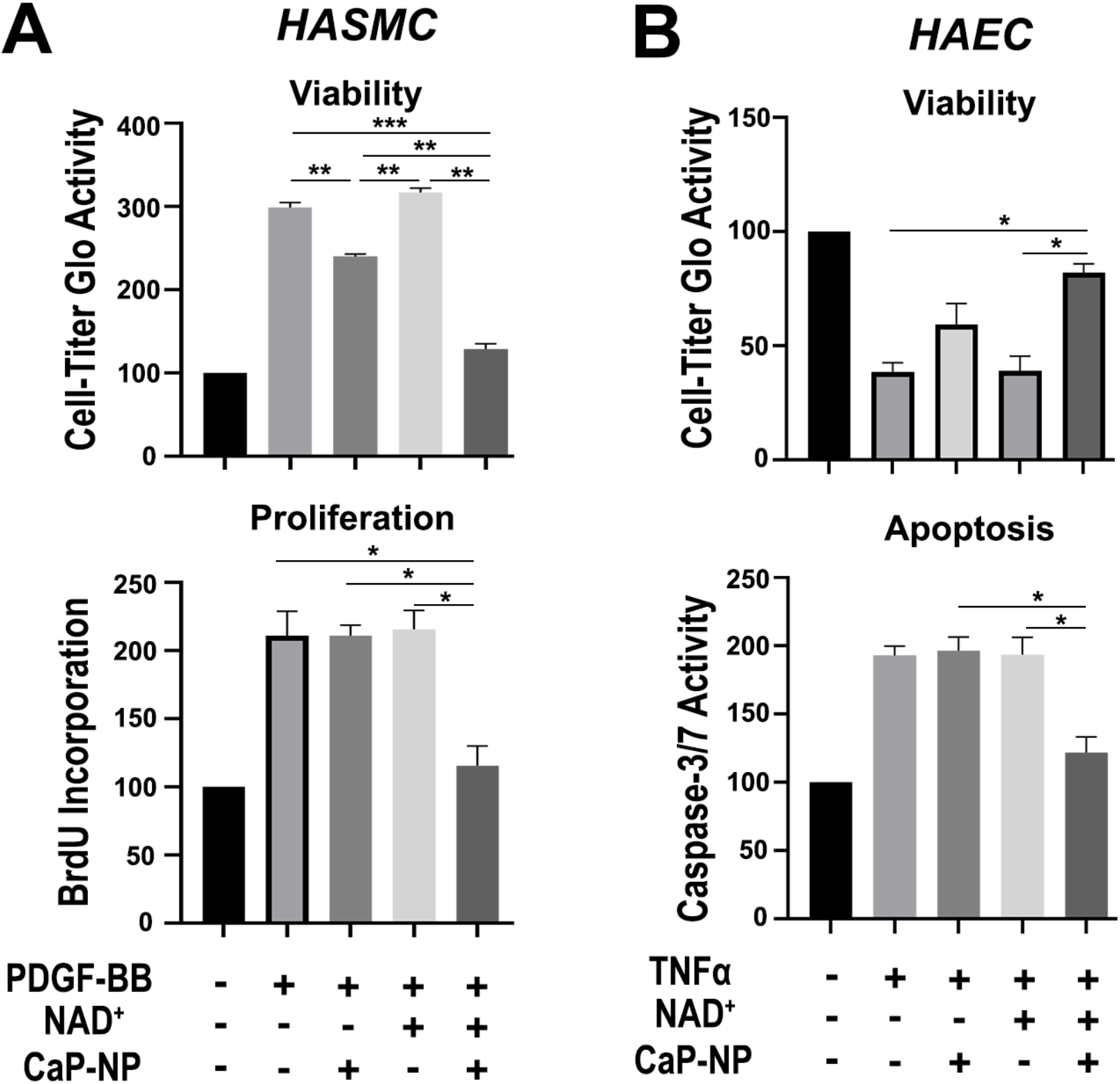
Platelet membrane-coated, NAD^+^-loaded CaP-NP (P-NAD^+^-NP) inhibits SMC proliferation while differentially preserving EC health. Primary culture of human aortic SMC (HASMC) and human aortic EC (HAEC) were utilized to test the differential cellular outcomes of Ca-NP-enabled NAD^+^ repletion. (A) Upon exposing 80-90% confluent and overnight starved HASMC to the respective treatments (equivalent to 10μM NAD^+^ payload) with or without 20 ng/mL PDGF-BB to stimulate SMC proliferation, cell viability and proliferation were determined using bioluminescent CellTiter-Glo Cell Viability assay (72hr) and colorimetric BrdU Incorporation assay (24hr). Presented as Mean ± SEM (n=4). (B) Upon exposing fully confluent HAEC to the respective treatments (equivalent to 10μM NAD^+^ payload) with or without 20 ng/mL TNFα to stimulate EC death, cell viability and apoptosis were determined using bioluminescent CellTiter-Glo Cell Viability assay (24hr) and Caspase3/7-Glo assay (4hr). Presented as Mean ± SEM (n=4). \**P* < 0.05, \*\**P* < 0.01, \*\*\**P* < 0.001, determined by One-way ANOVA followed by post-hoc Tukey test.

### 3.2 Biomimetic P-NAD^+^-NP Differentially Inhibits SMC Proliferation While Preserving EC Health

Prior reports^24^, albeit sparse, suggest a critical role of NAD^+^ biosynthesis (i.e., via the nicotinamide phosphoribosyltransferase (NAMPT)-mediated salvage pathway) in modulating the phenotypic transition of SMC under diseased states. Inspired by the outstanding intracellular uptake efficiency of the core CaP-NP as recently demonstrated^23^, we tested the *in vitro* efficacy of the platelet membrane-coated NP, with a special emphasis on probing the physiological impact of NAD^+^ repletion on SMC’s hyper-proliferative phenotypic transition. As shown in Figure 2A, the primary culture of human aortic SMC (HASMC) was stimulated with platelet-derived growth factor BB (PDGF-BB), a widely adopted stimulant of SMC proliferation that is abundant upon angioplasty as well as during the pathogenesis of restenosis^33^. Concomitant with PDGF-BB stimulation, SMC was also treated with either PBS, free NAD^+^ solution, empty biomimetic CaP-NP (empty NP), or biomimetic P-NAD^+^-NP (all at a concentration equivalent to 10μM NAD^+^ payload as determined in pilot studies). In stark contrast to the hyper-proliferative phenotype demonstrated in SMC treated with PDGF-BB, NAD^+^-replenished SMC displayed a quiescent phenotype following treatment with biomimetic P-NAD^+^-NP, as evidenced by significantly reduced cell viability and DNA synthesis (BrdU incorporation during the S phase of the cell cycle). Free NAD^+^ solution alone failed to inhibit SMC proliferation, as expected given its inability to bypass the plasma membrane. Similarly, empty NP was largely ineffective in modulating PDGF-BB-induced SMC hyper-proliferation. It is worth noting that a statistically significant, yet marginal benefit was indeed observed with the empty NP treatment group in terms of reducing SMC viability.

The benefits of NAD^+^ repletion in rescuing EC from programmed cell death have been documented, including in our recent report. Considering the lack of anti-restenotic therapies that could simultaneously promote EC regeneration, we hypothesized that our biomimetic P-NAD^+^-NP could offer a promising solution to this unmet medical need. As shown in Figure 2B, the primary culture of human aortic EC (HAEC) was challenged with tumor necrosis factor-alpha (TNFα) to induce cell death, mimicking the catastrophic cellular events following angioplasty-induced endothelium denudation. Similar to the experimental design in HASMC, TNFα-challenged EC was treated with biomimetic P-NAD^+^-NP (with NAD^+^ at 10 μM), as well as the appropriate control treatments. TNFα dramatically compromised EC viability and elicited apoptotic activation in HAEC, both of which were successfully rescued upon treatment with biomimetic P-NAD^+^-NP. No protective effect could be observed in all control groups, in which EC was treated with PBS, free NAD^+^, or empty NP.

### 3.3 Targetability, Biodistribution, and Biocompatibility of Biomimetic P-NAD^+^-NP

To investigate the IH lesion targetability of the platelet membrane-coated biomimetic CaP-NP, we loaded ATTO550-conjugated oligoRNA as a fluorescent tracer in the biomimetic P-NAD^+^-NP. Immediately following balloon angioplasty in the left common carotid arteries in male SD rats, the fluorescently labeled biomimetic P-NAD^+^-NP was intravenously injected immediately post procedure (at 10 mg/kg equivalent to the payload). At 6 hr post-injection, the injured carotid arteries, together with the contralateral un-injured carotid arteries as well as major organs, were collected for *ex vivo* fluorescent IVIS imaging. Consistent with prior efforts utilizing platelet membrane-coated NP of similar sizes but distinct composition^27,34,35^, our current study revealed higher enrichment of the biomimetic P-NAD^+^-NP in injured carotid arteries over uninjured ones (Figure 3B). Moreover, breaking away from earlier generations of biomimetic nanoplatforms, the platelet membrane-coated P-NAD^+^-NP displayed greater tropism toward arterial/aortic tissues over other major organs, especially the liver and spleen in the reticuloendothelial system, as demonstrated by the significantly greater level of normalized fluorescent signal as shown in Figure 3C. Overall, in line with our prior studies utilizing the platelet membrane coating for surface functionalization, the IVIS imaging analysis strongly supports the IH lesion-targetability and desired biodistribution pattern of the biomimetic P-NAD^+^-NP.

**Figure 3.**
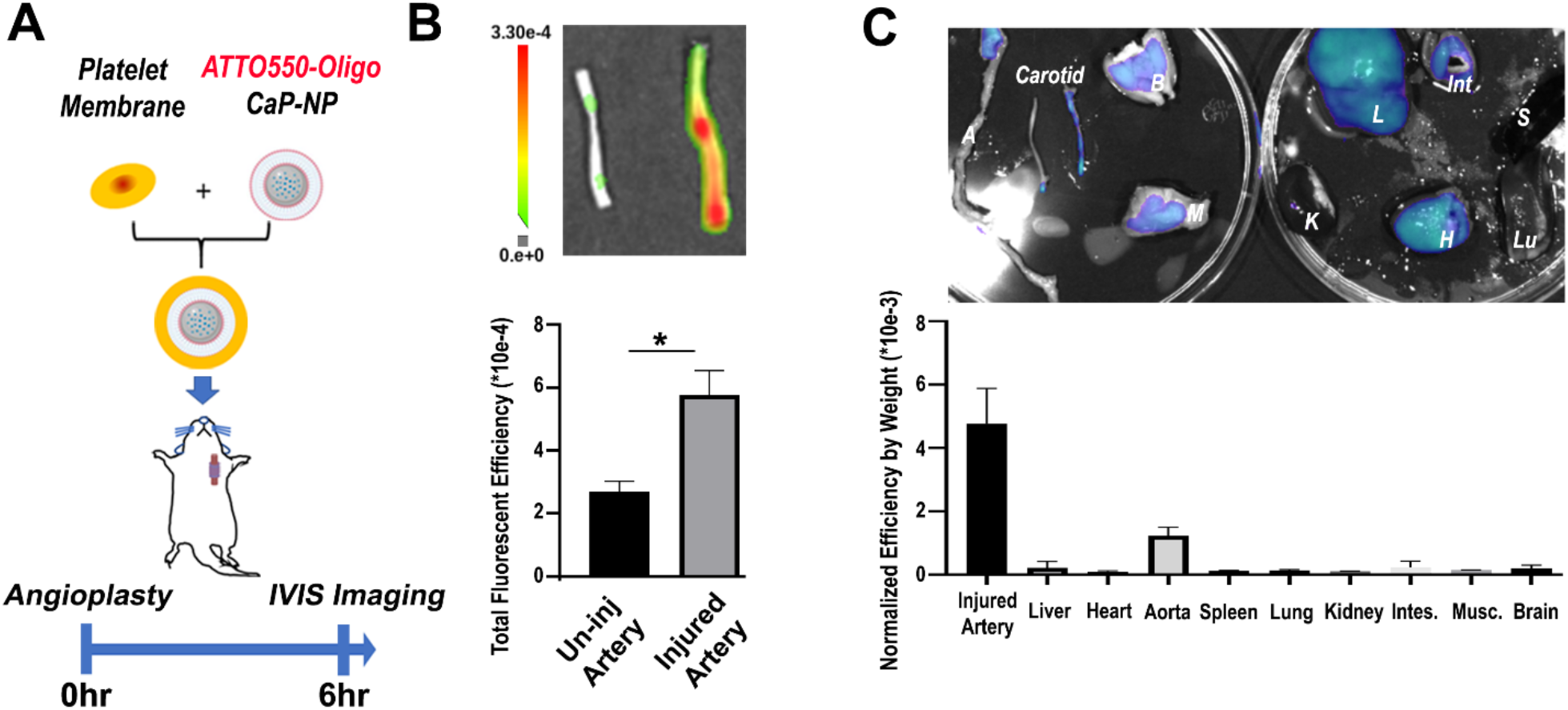
The platelet membrane-coated CaP-NP demonstrated selective lesion-targetability and desirable biodistribution. (A) Overall study design: ATTO550-oligoRNA-loaded biomimetic CaP-NP were intravenously injected immediately post-procedure. The balloon-injured and non-injured carotid arteries were collected 6hr post procedure/injection for *ex vivo* imaging using an IVIS system (Ex/Em: 545/575 nm). (B) *Ex vivo* fluorescence IVIS imaging of the injured (by balloon angioplasty) versus non-injured, contralateral carotid arteries. Quantitative analysis of the normalized fluorescence intensity was conducted. (C) *Ex vivo* IVIS imaging of injured carotid artery versus major organs. L, H, A, S, Lu, K, Int, M, and B represent liver, heart, aorta, spleen, lung, kidney, intestine (intes.), skeletal muscle (musc.), and brain. Quantitative analysis of the mean fluorescence intensity per unit mass in each organ or tissue shown in the *ex vivo* images.

Moreover, the biomimetic P-NAD^+^-NP is highly biocompatible in rodent models. In male SD rats that received a one-time injection of the biomimetic P-NAD^+^-NP (equivalent to 10 mg/kg NAD^+^ payload as established in our prior study), we observed no signs of toxicities and tissue structural damages in major organs including liver, spleen, heart, and kidney (Supplementary Figure 3), thereby supporting the feasibility of a more frequent and flexible injection regimen of the NAD^+^ repletion therapy.

### 3.4 Targeted NAD^+^ Delivery via the Biomimetic P-NAD^+^-NP Mitigates Post-angioplasty IH In Vivo

Inspired by our biomimetic P-NAD^+^-NP’s potent effect in inhibiting SMC hyper-proliferation and demonstrated targetability, we set to explore its anti-restenotic efficacy *in vivo*. Immediately following balloon angioplasty of left common carotid arteries in male SD rats, we provided a single-dose regimen via intravenous injection with the following treatments: saline, empty biomimetic CaP-NP (empty NP), free NAD^+^ solution (free NAD^+^), and biomimetic P-NAD^+^-NP — all at a dose equivalent to 10 mg/kg NAD^+^ payload. Rats were euthanized two weeks post angioplasty and treatment, followed by histological and morphometric analysis of carotid arteries to determine the lesional extent of IH. As shown in Figure 4B, accumulation of hyper-proliferative SMC could be observed in the neointimal layer in angioplastied carotid arteries. Further evaluation of IH, as determined by the intima-to-media ratio (I/M ratio), revealed the profound anti-restenotic benefits of NAD^+^-NP in comparison to the saline control group in Figure 4C. As expected, free NAD^+^ solution displayed no observable effects in altering the IH lesional size. Treatment with the empty NP, however, showed a marginal and statistically non-significant trend toward IH reduction, in accordance with the similar *in vitro* observation in reducing SMC viability as noted in Figure 2A. Analysis of the mRNA level of SMC contractility/maturation gene αSMA further corroborated the efficacy of the P-NAD^+^-NP in mitigating post-angioplasty IH (Figure 4D).

**Figure 4.**
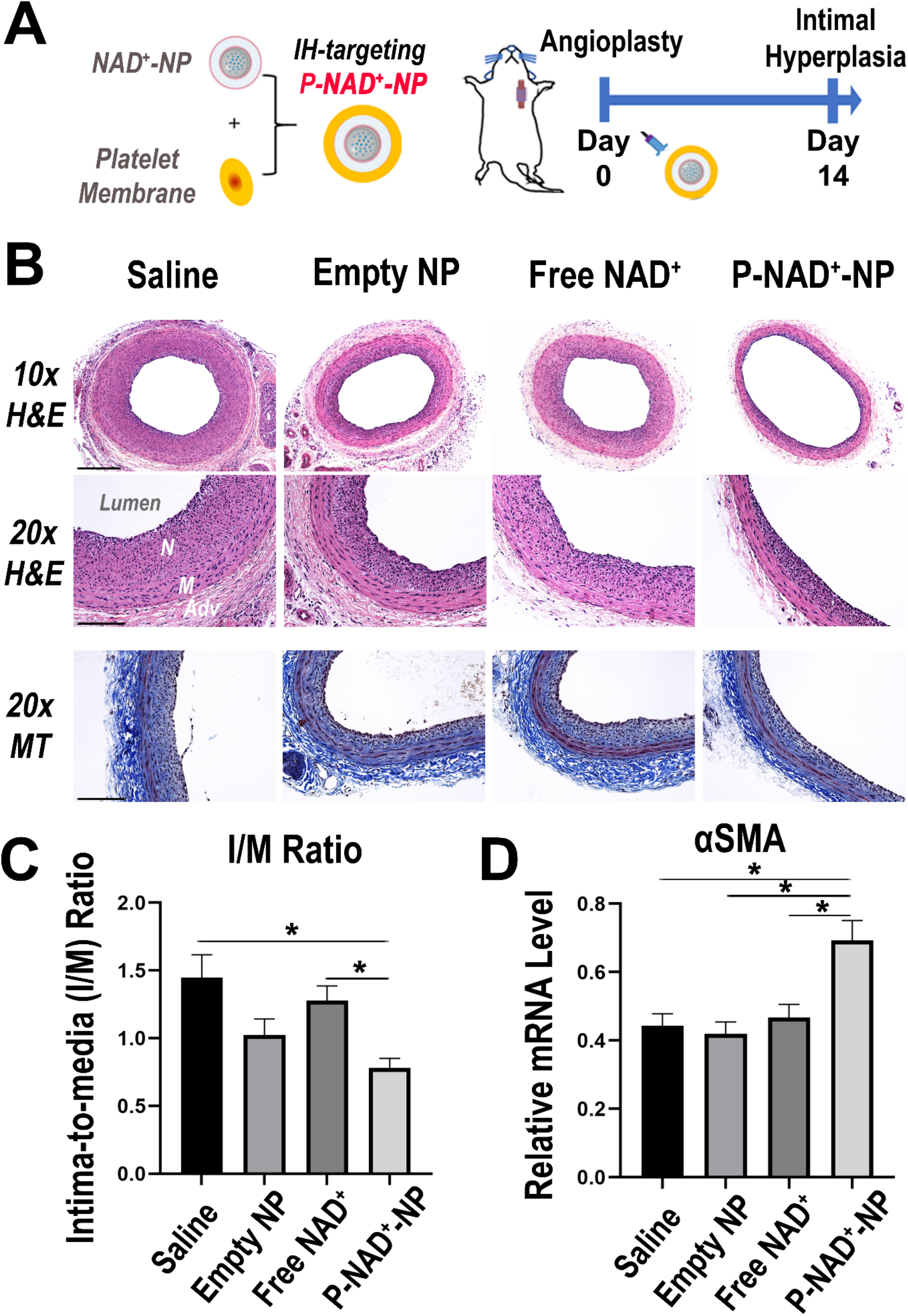
Targeted NAD^+^ delivery via the biomimetic P-NAD^+^-NP mitigated post-angioplasty IH in *vivo*. (A) Overall design: Prototypic NAD^+^-NP was coated with platelet membrane coating to enable IH-lesion targetability as well as biocompatibility. The injectable NAD^+^-NP or the control groups were intravenously administered (one-time injection; 10mg/kg NAD^+^ payload or equivalent) immediately following carotid artery balloon angioplasty in male SD rats. (B) Representative Hematoxylin and Eosin (H&E) and Masson’s Trichrome (MT) histology images of IH lesions (low and high magnitudes) at day 14 post-procedure. Neointima (N), media (M), and adventitia (Adv) are indicated, respectively. Scale bars: 275μm under 10x; 125μm under 20x. (C) Morphometric analysis of IH based on intima-to-media (I/M) ratio. (D) qPCR analysis of mRNA expression changes in tissue homogenates of rat carotid arteries post angioplasty. Quantitation of mRNA level of SMC contractility/maturation gene (αSMA). n=4-8. Presented as Mean±SEM. *P<0.05. One-way ANOVA with Tukey post-hoc analysis.

### 3.5 Targeted NAD^+^ Delivery via the Biomimetic P-NAD^+^-NP Accelerates Re-endothelialization Following Angioplasty-induced EC Denudation In Vivo

Based upon the EC *in vitro* data and anti-restenotic efficacy data *in vivo*, we hypothesized that effective NAD^+^ repletion, enabled by our novel biomimetic P-NAD^+^-NP may hold the key to solving the seemingly irreconcilable dilemma between inhibiting SMC proliferation and promoting EC regeneration. To test this, we utilized the same balloon angioplasty model in rat carotid arteries to induce endothelium denudation, immediately followed by a single-dose regimen of the P-NAD^+^-NP (equivalent to 10 mg/kg NAD^+^ payload) and the aforementioned control treatments. On Day 6 post angioplasty, animals were subjected to intravenous injection with Evans Blue dye to help determine the extent of re-endothelialization. As shown in the longitudinally opened carotid arteries in Figure 5B, the re-endothelialized luminal areas were not permeable to Evans Blue dye, whereas the non-EC-covered areas show prominent staining in blue color. Quantification of the re-endothelialization percentage over baseline level showed significantly improved endothelium regeneration following targeted NAD^+^ repletion with the biomimetic NAD^+^-NP over the control treatment group with saline. Similar to the observations in Figure 2A and Figure 4C, a slight yet statistically non-significant increase in re-endothelialization was noted following treatment with the empty NP. qPCR analysis of mRNA expression changes for genes representative of EC physiological function/integrity (eNOS), thrombogenicity (tissue factor/TF), impairment of EC healing (CXCL10), and inflammatory cytokine (ICAM1) further validated the EC protection of targeted NAD^+^ repletion *in vivo*. Taken together, our data strongly suggests that targeted NAD^+^ delivery, enabled by our innovative, biomimetic P-NAD^+^-NP, may offer a promising solution for safer yet effective management of restenosis.

**Figure 5.**
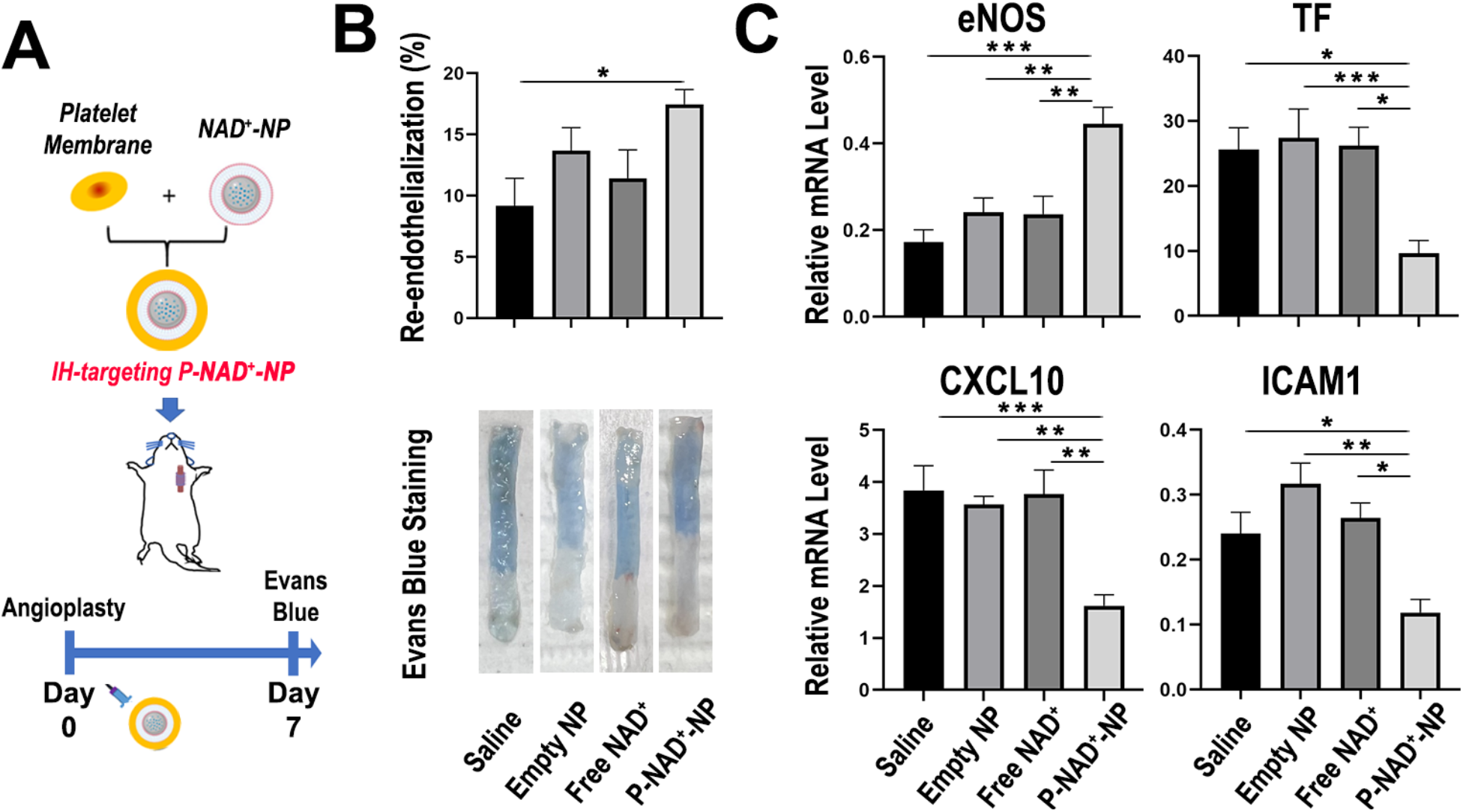
Targeted NAD^+^ delivery via the biomimetic P-NAD^+^-NP accelerated re-endothelialization following angioplasty-induced EC denudation *in vivo*. (A) The platelet membrane-coated NAD^+^-NP or the control groups were intravenously administered (one-time injection; 10mg/kg NAD^+^ payload or equivalent) immediately following carotid artery balloon angioplasty in male SD rats. Animals were injected with Evans Blue to visualize non-endothelium-covered luminal areas 30min prior to euthanasia and macroscopic examination at day 6 post angioplasty. (B) Representative images of Evans Blue-stained carotid arteries at day 6 post angioplasty and quantification of re-endothelialization (n=10-13). (C) qPCR analysis of mRNA expression changes in tissue homogenates of rat carotid arteries at day 6 post angioplasty. Quantitation of mRNA levels of EC signature gene representative of EC physiological function/integrity (eNOS), thrombogenicity (tissue factor/TF), impairment of EC healing (CXCL10), and inflammatory cytokine (ICAM1). Presented as Mean±SEM. *p<0.05. One-way ANOVA with Tukey post-hoc analysis.

## 4. Discussion

Despite decades of advancement in surgical techniques and device development, restenosis remains the primary cause of long-term failure following endovascular interventions for cardiovascular and peripheral vascular diseases.^1,2^ Numerous iterations of drug-eluting/coated endovascular devices have been widely adopted with demonstrated success in reducing the restenosis rate.^5,6^ Nevertheless, resolving treatment-resistant restenosis remains an unmet clinical need^36,37^; and in comparison to the lesions arising from bare metal stents and plain balloon angioplasties, the restenotic lesions following treatment with drug-eluting devices tend to be laden with significantly more neoatherosclerosis, hence posing a higher risk of severe complications.^38,39^ To make things worse, the often-toxic drugs used in balloon/stent have long been widely acknowledged to pose greater damage to the fragile EC than the phenotypically switched, hyper-proliferative SMC. Secondary to the damages caused by the drug and the drug-eluting devices, impaired EC regeneration following intervention — a process also known as re-endothelialization *in vivo* — predisposes the target vessel to a significantly higher risk of thrombosis and many more.^13-15^ Therefore, it remains a paramount challenge to develop new therapies that could discriminate against SMC proliferation hence preventing IH and restenosis, while preferentially preserving or even promoting EC regeneration. Although works from our group and many others have identified numerous disease-driving genes or pathways that are druggable or not, none have yet demonstrated clinical benefits nor a clear path toward clinical utility.

NAD^+^ is a key co-enzyme and metabolite involved in diverse biological processes such as bioenergetics, chromatin remodeling, redox homeostasis and immune responses.^40,41^ It impacts several signaling pathways implicated in the development and progression of vascular disease.^22^ Recent studies have highlighted the importance of the intracellular NAD^+^ pool in maintaining vascular homeostasis.^20,24,42^ NAD^+^ deficiency has been linked to EC dysfunction, aortic aneurysm, aortic stiffness, hypertension, and other vascular pathologies.^42-44^ The underlying mechanism behind NAD^+^’s role is multifaceted, including notable examples previously implicated in SMC/EC health and pathogenesis of IH, such as NAD^+^-dependent activation of sirtuin family (SIRTs) and deacetylation of substrate proteins, modulation of oxidative stress response, calcium signaling, and mitochondrial bioenergetics.^42,45-48^ Alterations in NAD^+^ biosynthesis and consumption pathways have also been suggested to contribute to the pathogenesis of cardiovascular and peripheral vascular diseases.^49-51^ Of particular note is NAMPT, the key enzyme responsible for the NAD^+^ salvage synthesis.^52^ Reduced NAMPT expression and concomitant NAD^+^ declination have been documented in HASMC in the context of aortic medial degeneration and aortic aneurysm^24^, in line with the widely reported pleiotropic role of NAMPT.^52^ However, recent evidence shows that its extracellular form, or eNAMPT, may play a disease-driving role in atherosclerosis, and similarly, in pulmonary SMC in the context of pulmonary arterial hypertension.^51,53,54^ While a growing body of evidence from our group and others supports the utility of NAD^+^ repletion in remedying pathological conditions involving endothelial and epithelial dysfunctions^23,55^, the functional link between NAD^+^ and aortic/arterial SMC/EC phenotypic modulation – especially in the context of restenosis and re-endothelialization – remains elusive. Our current study provides the first experimental evidence supporting the role of NAD^+^ repletion in the differential modulation of SMC’s proliferative phenotypic transition and EC’s apoptotic and inflammatory responses.

Strategies for boosting the intracellular NAD^+^ pool have demonstrated pleiotropic benefits in reversing aging and improving cardiometabolic processes^48,56^, but the path to clinical utility remains challenging due to the limited bioavailability and bioactivity of conventional NAD^+^ therapies.^57^ Cells cannot directly take up extracellular NAD^+^, which is negatively charged and susceptible to degradation.^58-60^ NAD^+^ has to be degraded extracellularly into its precursors, such as nicotinamide riboside (NR) and nicotinamide mononucleotide (NMN), which are subsequently taken up by cells and converted back to NAD^+^.^61^ Current NAD^+^ boosting strategies rely mainly on the supplementation of NAD^+^ precursors and other intermediates such as niacin, which need to be processed via salvage synthesis pathways which exhibit low speed and efficiency.^23^ While NAD^+^ precursors such as niacin have been previously explored as lipid-lowering agents^62^, and clinical efficacy in preventing IH and subsequent neo-atherosclerosis was not established.^63^ An emerging approach for boosting intracellular NAD^+^ is to activate NAD^+^ biosynthesis enzymes, but selective agonists remain at the early stages of preclinical evaluation.^64-66^ Collectively, these limitations necessitate an extremely high dose and treatment frequency, which greatly limit the clinical utility and efficacy of existing NAD^+^-boosting therapies.^67,68^ Recently, our team successfully achieved the first *in vivo* NAD^+^-replenishing therapy, built on an innovative nano-design featuring a lipid membrane coating and a calcium phosphate (CaP) core.^23^ The CaP-NP overcomes NAD^+^’s intrinsic limitations by protecting it from degradation and enabling direct intracellular NAD^+^ delivery through the endocytic pathway. Aside from the current study and our recent publication applying a non-targeted prototype in sepsis models^23^, there have been no prior *in vivo* studies using NAD^+^-loaded NP in any disease context, speaking to the paradigm-shifting nature of our technology. Compared with prior *in vivo* studies involving NAD+ repletion (or its precursors) that required highly frequent dosing at extremely high dosages (at least 100 mg/kg), our technology enabled outstanding preclinical efficacies at mere 1/10 or less dosage (10 mg/kg) with mere one dose over the course of 2 weeks.

The persistent lack of effective NAD^+^-boosting therapies testifies to the need for a new paradigm in NAD^+^ delivery. In the current study, we present the first effective NAD^+^-replenishing therapy for targeted NAD^+^ delivery to vascular lesions, built upon the technical innovations on two fronts — NAD^+^ nanoformulation and biomimetic surface functionalization. On the one hand, our prototypic NAD^+^-NP, without any specific targetability, conferred complete protection against endotoxin-induced EC dysfunction and mortality in three murine models of sepsis.^23^ Its lipid surface and CaP core afford a great extent of flexibility for further modification. On the other hand, as established by our study team, biomimetic surface modification with distinct cell membranes can grant NPs with lesion-targeting capacity and biocompatibility, which can be readily achieved on top of the prototypic NAD^+^-NP.^25^ Owing to the platelet’s “first responder” role by adhering to the exposed base membrane and extracellular matrix via its repertoire of membrane proteins (e.g., glycoprotein VI)^32^, the extracted platelet membrane coating, ergo the biomimetic P-NAD^+^-NP, could inherit such lesion-responding capacity. The abundant presence of immune modulatory cues on platelet membrane surfaces, such as CD47, also confers biocompatibility and reduced reticuloendothelial clearance, as demonstrated in our current and prior studies.

There are several limitations to the present study. Firstly, the empty NP displayed a certain level of benefits, albeit mostly numerical and non-significant, in mitigating SMC proliferation and IH while protecting EC recovery. Although future studies are still warranted to validate the potential of the “carrier” itself, we contend this unintended/additive advantage of the biomimetic P-NAD^+^-NP could lend further support to its translation to clinical utility. Secondly, our *in vivo* data were derived from otherwise healthy SD rats at a relatively acute endpoint (2 weeks post angioplasty). In real-world applications, however, the majority of patients requiring interventions are of diabetes and other comorbidities, with IH and restenosis progressively developing over the course of months to years. Therefore, any therapeutic strategy should not only be applied/tested to chronic IH lesions, but also in subjects with metabolic disorders (e.g., diabetic or dyslipidemic rats/mice). Future studies are needed to thoroughly test the translational potential of the targeted NAD^+^ repletion therapy in appropriate *in vivo* models that address these clinically relevant variables. Last but not least, while we observed remarkable *in vivo* efficacies in mitigating IH and promoting re-endothelialization with a mere one dose of targeted NAD^+^ repletion, it is possible that a more frequent injection schedule (e.g., weekly) of the biomimetic P-NAD^+^-NP — which demonstrated a high level of biocompatibility — could yield greater therapeutic outcomes.

## 5. Conclusions

The current standard of care for restenosis is far from optimal and is detrimental to the vascular endothelium. Our current study provides the proof-of-concept for targeted NAD^+^ repletion as a viable strategy for endothelium-friendly, anti-restenotic therapy. Built upon the combined merits of biomimetic surface functionalization and a first-in-class P-NAD^+^-NP, a platelet membrane-coated CaP-NP design was developed to achieve IH lesion-targeted, efficient intracellular NAD^+^ repletion. A one-time administration of biomimetic P-NAD^+^-NP sufficed to mitigate SMC proliferation and IH; in stark contrast to the status quo that impairs endothelium recovery, targeted NAD^+^ repletion rather promoted re-endothelialization and blocked EC inflammatory/thrombogenic phenotype of EC. Collectively, our biomimetic P-NAD^+^-NP may offer an alternative path toward the development of the next-generation, endothelium-protective, anti-restenotic therapy.

## Acknowledgments

This work was partially supported by the National Institute of Health (NIH) grants R01HL143469, R01HL129785, and R21AI165977.

