## Supplementary Information for "Targeted NAD^+^ Delivery for Intimal Hyperplasia and Re-endothelialization: A Novel Anti-restenotic Therapy Approach"

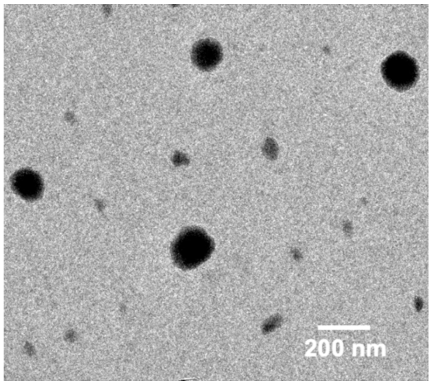


Supplemental Figure 1. TEM image of $NAD^+$-NP without platelet membrane coating.

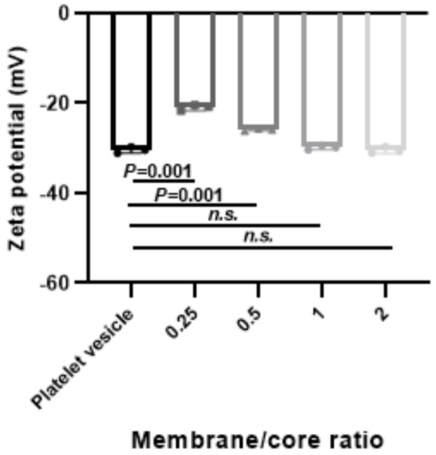


Supplementary Figure 2. Optimization of platelet membrane/$NAD^+$-NP ratio. Zeta potentials of platelet-derived vesicles and $NAD^+$-NP with different platelet membrane to $NAD^+$-NP weight ratios were measured. Data are presented as mean ± s.d. (n=3). Statistical significance was calculated by one-way analysis of variance (ANOVA) with Tukey's post hoc test. n.s., no significance (i.e., $P \geq 0.05$).

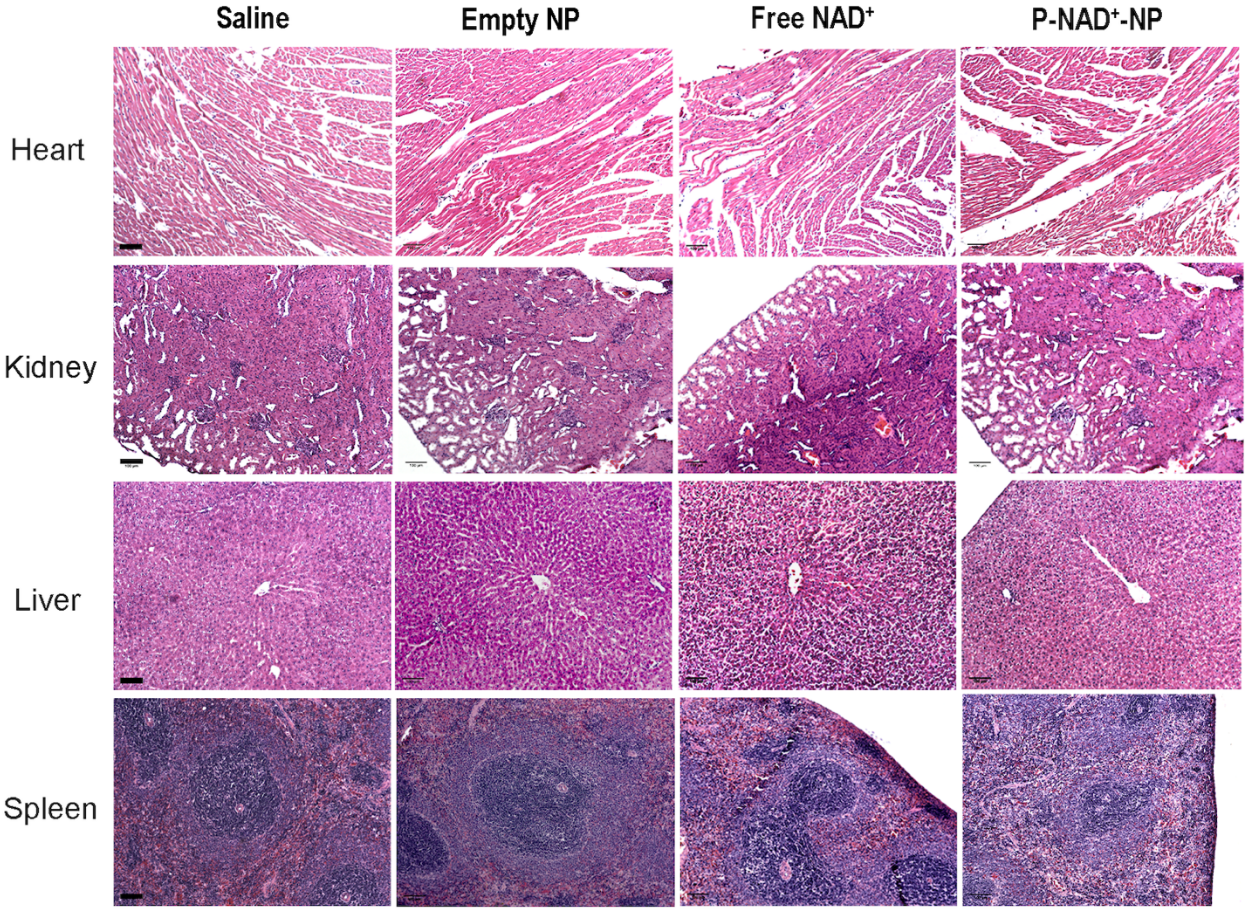

Supplementary Figure 3. Lack of systemic toxicity in major organs following biomimetic P-NAD$^+$-NP administration. Representative H&E images of heart, kidney, liver, and spleen from SD rats euthanized at day 14 post angioplasty and a single-dose regimen of P-NAD$^+$-NP and respective control treatments (equivalent to 10mg/kg payload). Scale bar: 100μm.

| Target Gene | Sequence |
|---|---|
| Rat αSMA | Forward: ACCTTCAATGTCCCTGCCATGTA |
| | Reverse: ACGAAGGAATAGCCACGCTCA |
| Rat eNOS | Forward: GAGCCCCCAGAACTCTTCAC |
| | Reverse: CAATGGTCACTTTGGCCAGC |
| Rat Tissue Factor | Forward: TTGGAGTGGCAACCGAAACC |
| | Reverse: ACAATCTCGTCGGTGAGGTC |
| Rat CXCL10 | Forward: TGCAAGTCTATCCTGTCCGC |
| | Reverse: TGACCTTCTTTGGCTCACCG |
| Rat ICAM1 | Forward: AGGATCGCTGCTCGGAAAAT |
| | Reverse: GCGGGGTATATGGTGTCAG |
| Rat GAPDH | Forward: AAGGTCGGTGTGAACGGATTT |
| | Reverse: CTTTGTCACAAGAGAAGGCAGC |

Supplementary Table 1. Primer sequences for qPCR assay
